# Mutations in coral soma and sperm imply lifelong stem cell renewal and cell lineage selection

**DOI:** 10.1101/2021.07.20.453148

**Authors:** Elora H. López-Nandam, Rebecca Albright, Erik A. Hanson, Elizabeth A. Sheets, Stephen R. Palumbi

**Affiliations:** Biology Department, Hopkins Marine Station of Stanford University, Pacific Grove, California 93950 USA; Institute for Biodiversity and Sustainability Science, California Academy of Sciences, San Francisco, California, 94118, USA

## Abstract

In many animals, the germline differentiates early in embryogenesis, so only mutations that accumulate in germ cells are inherited by offspring^1^. Exceptions to this developmental process may indicate that other mechanisms have evolved to limit the effects of deleterious mutation accumulation^2^. Stony corals are animals that can live for hundreds of years^3^ and have long been thought to produce gametes from somatic tissue^4^. To clarify conflicting evidence about germline-soma distinction in corals, we sequenced high coverage, full genomes with technical replicates for parent coral branches and their sperm pools. We identified post-embryonic single nucleotide variants (SNVs) unique to each parent branch, then checked if each SNV was shared by the respective sperm pool: 26% of post-embryonic SNVs were shared by the sperm but 74% were not. We also identified germline SNVs, those that were present in the sperm but not in the parent. These data suggest that self-renewing stem cells in corals differentiate into germ and soma throughout the adult life of the colony, with SNV rates and patterns differing markedly in stem, soma, and germ lineages. In addition to informing the important place in the evolutionary spectrum from non-Weismannian to Weismmanian animals that corals occupy, these insights inform how corals may generate adaptive diversity necessary in the face of global climate change.

## Main

In 1889, Weismann proposed that germ and somatic tissues serve starkly different functions: germ cells protect heritable information and pass it on to the next generation, while somatic cells perform the functions necessary to keep an organism alive but do not contribute to the genetic makeup of the organism’s offspring. This would explain why mutations that accumulate in somatic tissues during an animal’s lifetime– including those that cause cancer-are not inherited by that animal’s offspring. Instead, only mutations in germ cells, which undergo fewer cell divisions and have lower mutation rates, are inherited^5,6^. Since Weismann, embryonic germ-soma separation has been shown in vertebrates and many other animal taxa, but not in plants or in some animal groups, including cnidarians, sponges, tunicates, and platyheminths^7,8^.

Potential animal exceptions to Weismann’s Germ Plasm Theory are intriguing because they may have novel mechanisms to reduce the number of deleterious mutations inherited by sexually produced offspring. Moreover, such exceptions may signal the potential existence of stem cell lineage types not seen in vertebrates. For example, the model cnidarians *Hydra* and *Hydractinia* possess interstitial stem cells, denoted i-cells, that can differentiate into both germ and soma during adult life^9,10^. A few models have hypothesized how heritable post-embryonic mutations may affect the gamete pool^11–13^, but there are very few datasets on the pattern of somatic mutations and their inheritance in long-lived animals^14,15^.

Clonal, colonial corals can live for hundreds to thousands of years, and were long thought to generate gametes from the somatic cells of clonal polyps^4^. Coral colonies accumulate somatic mutations at a rate similar to noncancerous human tissues^14^. If these mutations are inherited by the coral’s gametes, they may increase the heritable mutational load of these animals. Some previous studies identified putative somatic mutations in the gametes or juvenile offspring of mutant parents^16,17^, but others have reported absence of somatic mutations in the gametes^18^. These studies tracked few mutations, ranging from 9 to 170, and none detected germline mutations in gametes or offspring. Only one verified that their putative mutations were not PCR or sequencing error^18^. Here, we interrogated, by sequencing full genomes and verifying with technical replicates, multiple branches from multiple coral colonies and their sperm. We identified germline variants in the sperm as well as post-embryonic variants in the parent. The data reject the hypothesis that somatic cells give rise to germ cells in corals, but also reject the hypothesis that corals possess embryonic germline differentiation. Rather, we show that both parent tissue and sperm arise from a common stem cell lineage that proliferates and differentiates throughout the long lives of these animals. Our data indicate **not** that somatic mutations are inherited by germ cells, but that mutations identified in both soma and germ cells are a result of inheritance from a common progenitor stem cell lineage. This is a significant departure from what the two previous hypotheses were (either that somatic mutations get inherited by the sperm, or that the development of germline ends at the embryonic stage).

## Results

To clarify the inheritance of mutations and the presence of germline-soma distinction in *Acropora hyacinthus*, we removed branches from soon-to-spawn adult coral colonies and placed them into individual cups of seawater (Extended Data Fig. 1). Each branch released gamete bundles into its respective cup 20 minutes later (Fig 1, Extended Data Fig. 1b). We extracted DNA from each branch and each sperm pool, then constructed two replicate full genome libraries from each DNA extraction (Fig 1b). To be verified, a SNV had to be present in both replicate libraries of a given sample. The technical replicates eliminated over 90% of putative SNVs that would have been called if we had used one library per sample, although the exact number varied by SNV category (Extended Data Figs. 2, 3).

**Figure 1.**
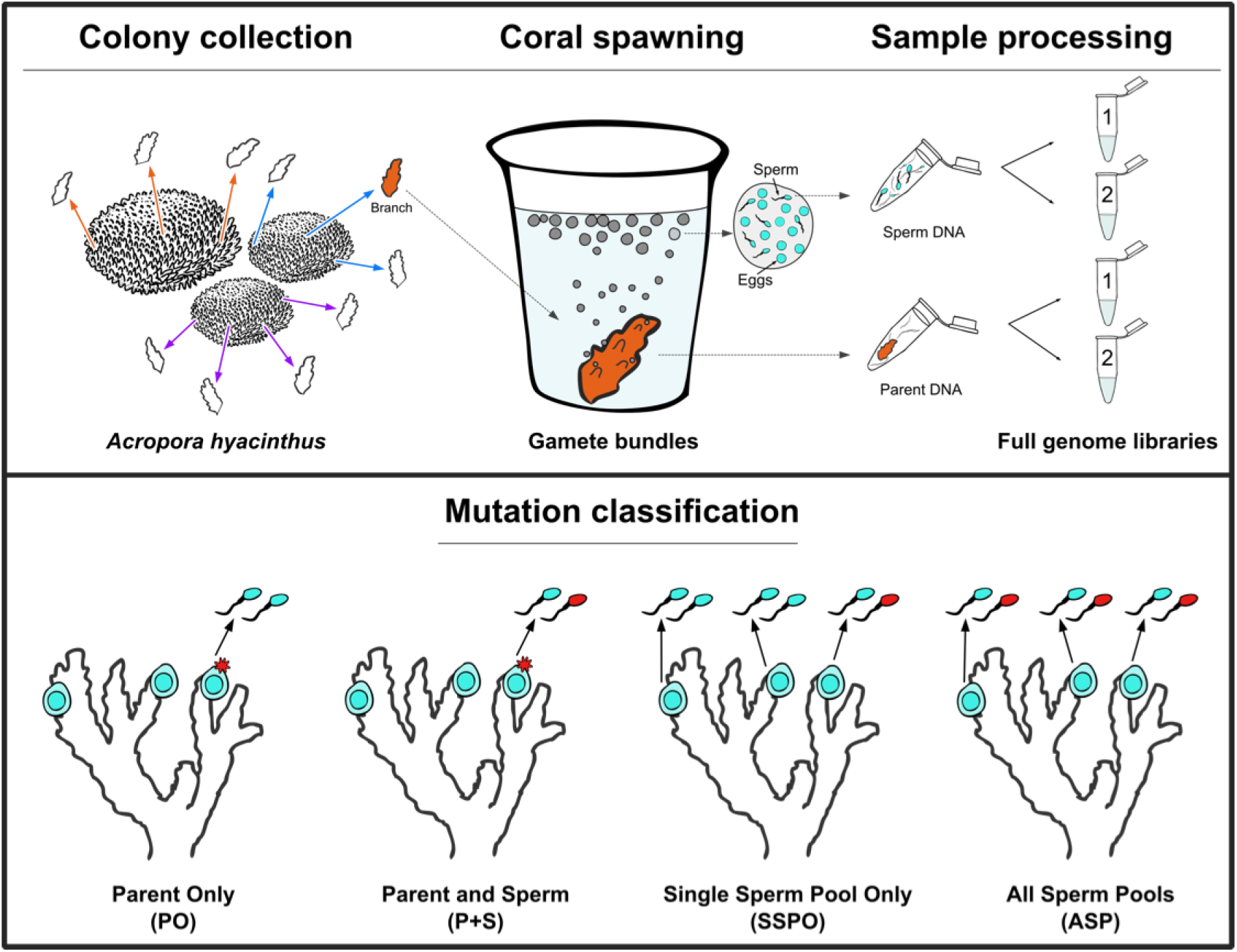
Data collection (top) and mutation classification (bottom). Top: Twenty minutes prior to spawning, 3-4 branches were broken off of 3 parent colonies and placed into individual cups of seawater, for a total of 10 branches in cups. Branches then released eggs and sperm into each cup, and sperm was collected from the cup. Both the sperm pool and the parent samples were stored in RNAlater and frozen. Genomic DNA was extracted from each parent branch and sperm pool (see Methods). For each genomic DNA extraction we constructed two full genome libraries (see Methods) for technical replication. Mutation classifications (bottom, left to right): 1.) A mutation unique to a single branch of the colony, but the sperm from the branch does not share the mutant genotype (Parent Only, PO). 2.) A mutation unique to a single branch of the colony, and the sperm from the branch shares the mutant genotype (Parent and Sperm, P+S). 3.) A mutation unique to just one sperm pool in the colony, not shared by other sperm pools or the parent branches (Single Sperm Pool Only, SSPO). 4.) A mutant genotype shared by all sperm pools from a particular colony, but none of the parent branches in that colony (All Sperm Pools, ASP). Figure by Shayle Matsuda.

We identified four different types of SNVs: those that were unique to the polyps from a single parent branch in a colony but were not detected in the sperm from that branch (Parent Only) (1c), those that were found in just a single parent branch in a colony and were also shared by the sperm from that branch (Parent and Sperm) (1d), those that were unique to a single sperm pool in a colony and not present in any branch of the colony (Single Sperm Pool) (1e), and those that were shared by all sperm pools in a colony but had never been seen in the polyps from any branch (All Sperm Pools) (1f).

We assayed nine parent polyp samples, and the respective sperm pools for seven of those samples, across three different colonies. The average depth of coverage across the genome was 40.6 ±3.1 (1 s.e.m) for the parent polyp libraries and 65.2 ±6.9 (1 s.e.m) for the sperm pool libraries (Supp. Table 1). Across the full dataset we identified 2,356 SNVs. All but one of these SNVs were at unique sites, indicating that the SNVs called were not a result of consistent mapping error or bias (Supp. Table 2). Each SNV was classified as a Gain of Heterozygosity (GoH), in which the aberrant sample was a new heterozygote and all others were homozygous, or a Loss of Heterozygosity (LoH), in which the aberrant sample was homozygous and the other samples were heterozygous.

We identified 146-351 post-embryonic SNVs per parent branch (Supp. Table 1). The rate of SNVs unique to a single parent branch, but not found its sperm pool (Parent Only, PO) ranged from 0.90-2.55 x 10^-6^ SNVs/bp per branch, with an average rate of 1.76 ± 0.23 x10^-6^ (1 s.e.m.) SNVs/bp per branch (Figure 2a). The rate of SNVs unique to a single parent branch, and shared by its respective sperm pool (Parent and Sperm, P+S) ranged from 0.34-0.96 x10^-6^ SNVs/bp per branch, with an average rate of 0.59 ± 0.1 x 10^-6^ (1 s.e.m.) SNVs/bp per branch (Figure 2a). On average, the rate of Parent Only SNVs was 3.4 times higher than the rate of Parent and Sperm SNVs found in a given branch. On average, 25.7% ±3.7% (1 s.e.m.) of SNVs found in a parent branch were Parent and Sperm SNVs, while 74.3% ±3.7% (1 s.e.m.) post-embryonic SNVs in a branch were Parent Only SNVs. These findings contradict the hypothesis from the Germ Plasm Theory that post-embryonic mutations would not be found in the sperm at all. They also contradict the common assumption that all coral somatic cells can produce gametes.

**Figure 2.**
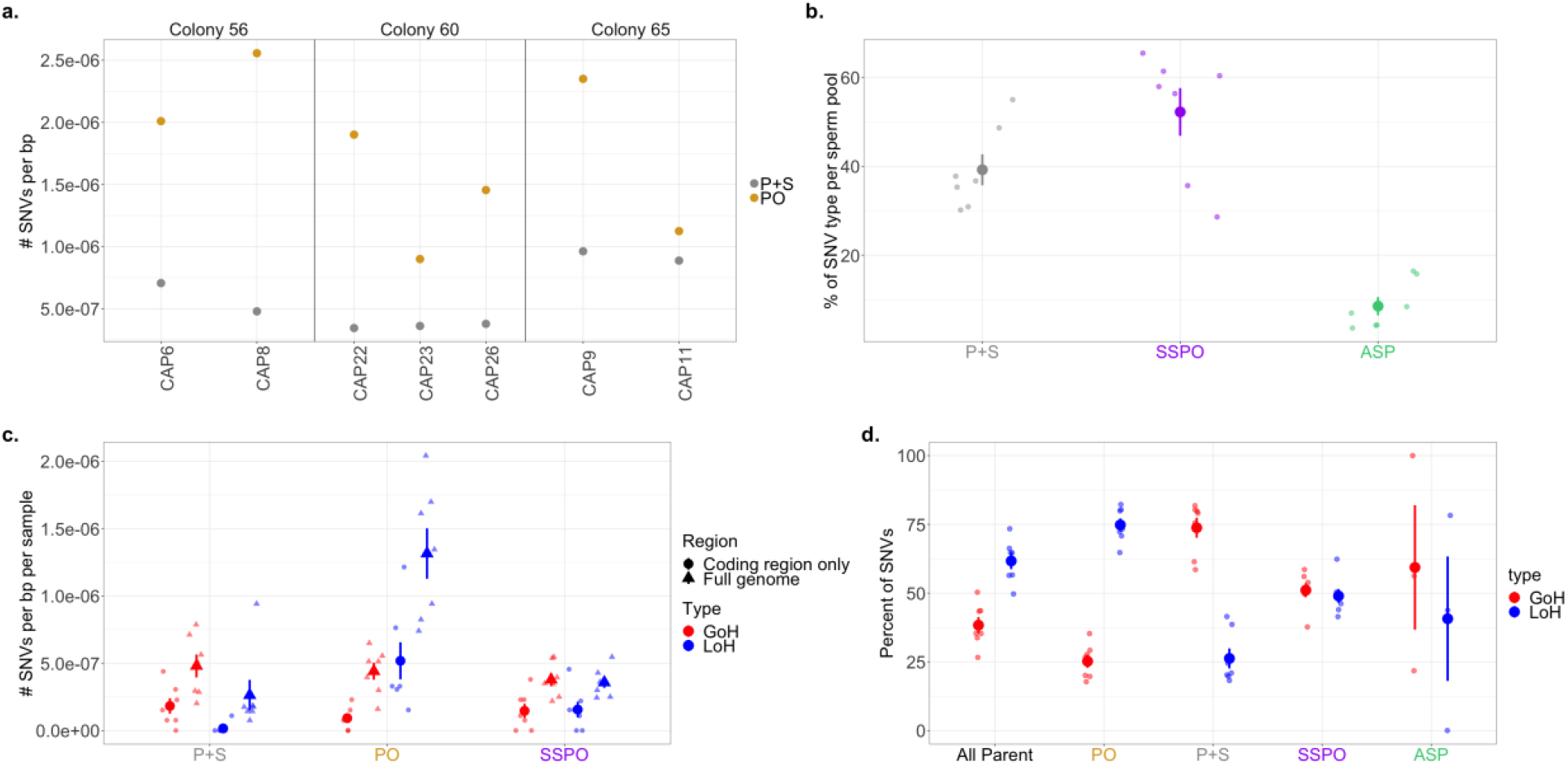
SNV rates and proportions across different classifications. a.) The rate of SNVs per bp for two SNV types: shared by parent and sperm (gray) and found in the parent only (yellow) for the seven parent-sperm pairs from the three colonies. b.) The average percentage of SNV type (parent and sperm, P+S), (parent only, PO), and all sperm pools (ASP) found in each sperm pool sample (N=7). c.) The average rate of SNVs per bp per sample (N=7) across the full genome and for the coding regions only, for three SNV types: parent and sperm (P+S), parent only (PO), and single sperm pool only (SSPO). For each SNV type, each subtype (GoH and LoH). d.) The average percentage of SNVs that were GoH and LoH for each of the four SNV types found in each sample (N=7). For b, c, and d, the mean for each category is shown as a large point with error bars extending out; error bars represent ±1 s.e.m. Each individual data point (N=7 for P+S, SSPO, PO, and All Parent, N=3 for ASP) is shown as a smaller point for each category.

Of the post-embryonic SNVs found in the sperm pools, 39.2% ±3.5% were Parent and Sperm (Fig. 2b), the same Parent and Sperm SNVs as shown in Fig. 2a. An additional 52.2% ±5.4% of SNVs in the sperm pools were found only in the sperm pool (Single Sperm Pool Only, SSPO) (Fig. 2b). A small number of sperm SNVs, 8.5% ±2.1%, were found in all sperm pools from that colony but none of the parent samples that spawned them (All Sperm Pools, labelled ASP, Fig. 2b). That 2 out of every 5 SNVs present in a given sperm pool are post-embryonic, non-germline variants indicates that the lack of an embryonic germline increases the number of SNVs in a colony’s gametes by 66%, compared to what the diversity would have been if the germline were segregated at the embryonic stage. This may help explain the high degree of heterozygosity in many stony coral species, though it is not yet known what fraction of these SNVs are too deleterious to survive into adulthood.

The rate of SNVs per bp was significantly higher across the full genome than the rate of SNVs in the coding regions of the genome for all SNV types (Fig. 2c) (see Table 1 for all means and Wilcoxon signed-rank test results). This may indicate that there is stronger purifying selection against SNVs in coding regions than in non-coding regions of the genome, or it may be a result of higher mismatch repair in exons^19^.

We examined the spectrum of mutations, the relative proportions of mutations in different classes, and found no significant differences in spectra among parent only, shared, and germ line specific mutations (Extended Data Fig. 5). These data confirm lack of a signature of UV-associated mutations in corals^13^, which is intriguing considering that these colonies grow in high UV conditions, and in highly oxygenated warm water^20^.

Losses of heterozygosity tend to arise as a result of gene conversion due to homologous recombination, a form of double stranded DNA break repair^21^. Consistent with previous findings^14^, 38.3 % ±3.0 % (1 s.e.m.) of all parent SNVs being GoH and 61.7 ±3.0% (1 s.e.m.) being LoH. SNVs that were shared by the parent branch tissues and the sperm had a much higher fraction of GoH and lower fraction of LoH (73.8 ±3.6% and 26.2 ±3.6%, respectively) than did parent SNVs that were not found in the sperm (25.2 ±2.4% and 74.8 ±2.4%, respectively) (Wilcoxon signed-rank test, V=28, p = 0.015) (Fig. 2d). SNVs found in just a single sperm pool had approximately equal proportions of each type, 51.1% ±2.5% GOH and 48.9 ±2.5% LOH SNVs. High LOH in soma that is not inherited by the sperm could be due to high incidence of double-strand breaks in somatic cells exposed to high light and photosynthetically derived oxidation, or high LOH levels in the Parent Only SNVs may reflect stronger selection against GOH than LOH in differentiated somatic cells.

To explore the role of selection on patterns of genome change, we compared the rates of missense and synonymous SNVs across four classes: all somatic SNVs, Parent Only SNVs, Parent and Sperm SNVs, and Single Sperm Pool Only SNVs. There were no coding SNVs in the All Sperm Pool category. The average rate of coding mutations was highest in parent only SNVs (6.0 x10^-7^ ±1.6 x 10^-7^, Extended Data Fig. 6). Across the full *Acropora hyacinthus* proteome, we estimate that 78.1% of sites are nonsynonymous and 21.9% of sites are synonymous (Fig. 3). The percent of coding mutations that were missense was higher in Single Sperm Pool Only SNVs (73.7 ±11.8%) than in the other categories (55.2 ±6.8% all somatic, 51.5 ±10.7% Parent Only, 47.6 ±14.6% Parent and Sperm, Fig. 3). The higher mean percent missense in Single Sperm Pool Only SNVs was not statistically significant, likely attributable to the small number of coding mutations in each category. Like most studies on somatic mutations to date, the small number of coding mutations in this study (94) leaves us underpowered to detect selection^22^. However, the fairly consistent pattern of more missense mutations in sperm pool samples than somatic samples provides a first hint that the SNVs in the soma may experience stronger negative selection than germline SNVs.

**Figure 3.**
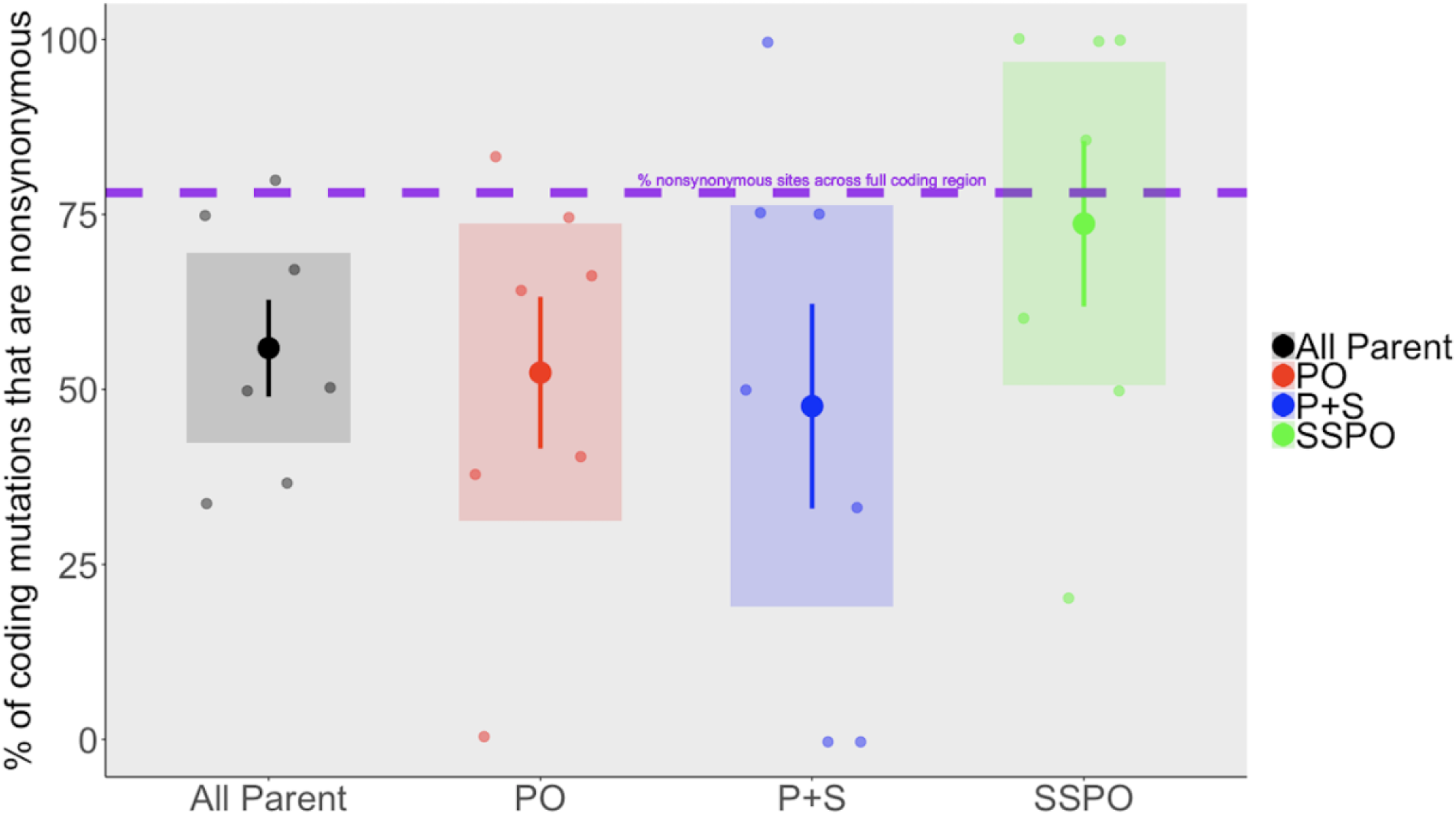
The mean percent of protein-coding SNVs that are nonsynonymous across coral branches (N=7) for All Parent, Parent Only (PO), Parent and Sperm (P+S), and Single Sperm Pool Only (SSPO) categories. Large circles indicate the mean, with error bars indicating ± 1 s.e.m. and shaded boxes indicating 95% CI. Each coral branch (N=7) is shown as a smaller point for each category.

## Discussion

Our data show that 26% of the SNVs that we identified in parent tissue were also present in the sperm spawned from that branch, and 74% were not. If we had found only separate parent and sperm SNVs this would have shown that *Acropora* corals have classical Weismannian germ and somatic cell lineage differentiation at the embryonic stage, which has been suggested previously^18^. Likewise, if we had found that all parent tissue SNVs were also in the sperm, we would have concluded that *Acropora* corals developed gametes directly from those tissues^16,17^. Because we found some SNVs shared by sperm but the majority not, we hypothesize that in colonial corals, shared parent and sperm SNVs derive from mutations in a common ancestor stem cell lineage that self-renews and proliferates through the colony, and that eventually differentiates into both germ and soma throughout the colony’s adult life. This type of lineage (i-cells) has been described in Hydrozoan cnidarians^9,10^. Although i-cells have not yet been identified in corals, cells that look like the i-cells are present in larvae of the coral *Acropora millepora*^23^. Our data suggest that branch-specific SNVs shared in germ and somatic cells first arose in an i-cell lineage proliferating in that branch, and then differentiated into germ and soma (Fig. 4). SNVs found only in the parent but not in the sperm would have arisen in terminally-differentiated somatic cells that cannot produce gametes, and SNVs found only in the sperm would have arisen from differentiated germ cells (Fig. 4).

**Figure 4.**
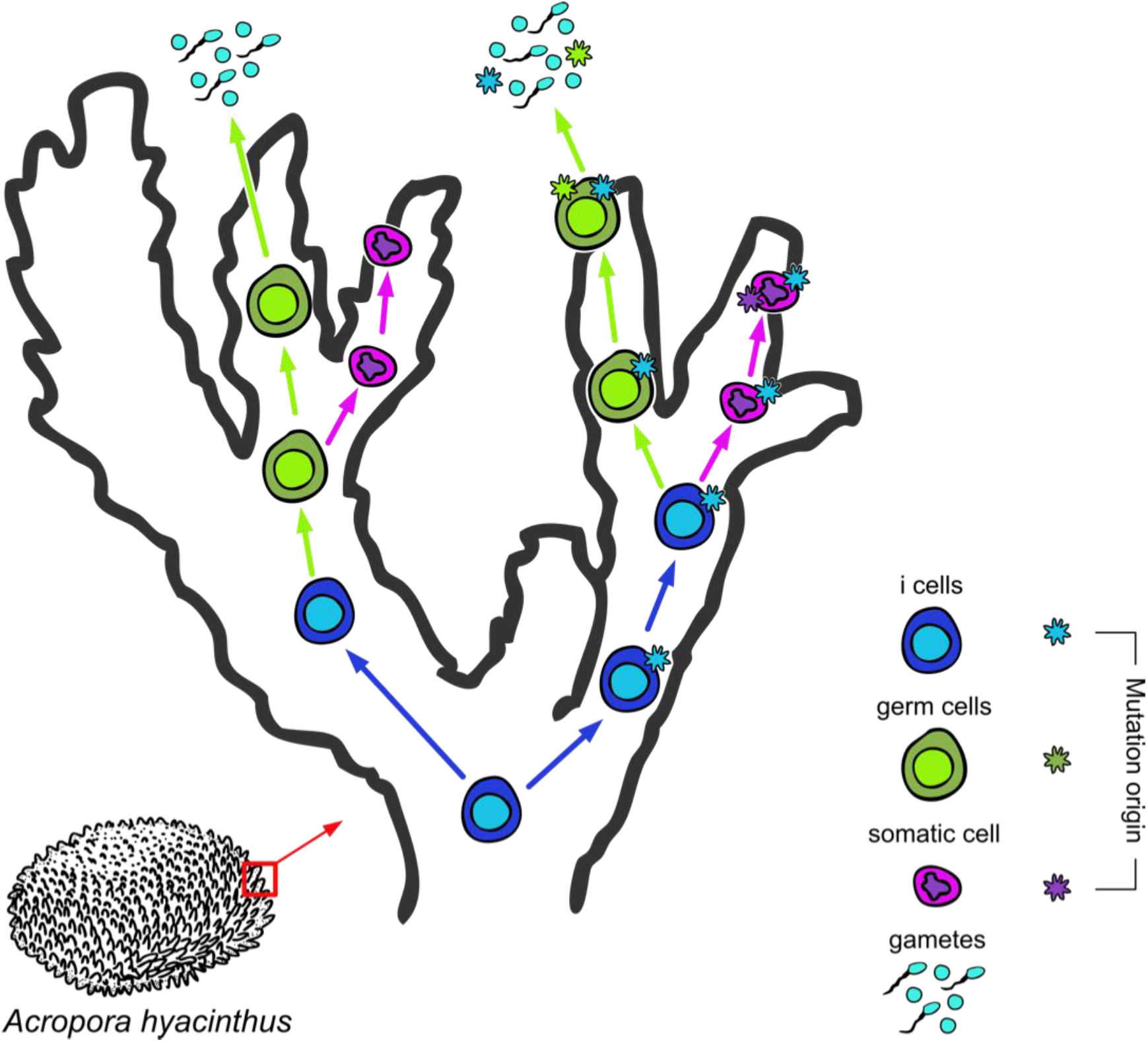
Schematic for how SNVs that arise at different stages of cell lineage development and differentiation proliferate. SNVs that arise in i-cell may be found in both germ and somatic cells later, if the mutant i-cell lineage differentiates into both germ and soma. SNVs that arise in soma or germ cell lineage post-differentiation will only be found in those differentiated lineages. Figure by Shayle Matsuda.

We hypothesize that the program of sequential germ line differentiation during adult life shown in Hydrozoans is likely a conserved trait across Cnidaria and was present in the cnidarian common ancestor. In corals, germ and soma differentiation appears to happen locally, resulting in evident mosaicism in every branch (Fig. 2a, Fig. 4). Solana^8^ made a similar prediction for planarians– when stem cells can differentiate into both soma and germ cells, mutations that appear in both the soma and the germline are derived from mutations in those stem cells.

Because the progenitor stem cell mutations are inherited by the soma, they are tested by natural selection/the environment in a way that the germ cell-derived mutations are not. For instance, if a stem cell mutates and then that mutant stem cell line gives rise to both soma and sperm, then that mutation is exposed to the environment in the form of tentacle, gastroderm, or other somatic cell types. If the mutation is deleterious for the soma of the polyp, then the polyp will be less likely to reproduce and pass on that deleterious mutation. If the mutation is neutral or advantageous for the polyp soma, then the polyp may survive to pass on the mutation, because that mutation is also in the polyp’s gametes. Note that this does not mean that there is inheritance from soma to sperm in this scenario (Fig. 4). Rather, the presence of the stem cell-derived mutations in the soma allowed the polyp to reproduce (or not), and because that stem cell-derived mutation was also in the gametes of the polyp, it can be inherited. This is a significant departure from the conclusions that other recent empirical papers have drawn, but it is very much in line with classical theory.

In support of the above hypothesis, we have evidence for some purifying selection on SNVs in stem and somatic cells, but not in germ cells. Germline coding SNVs showed the same rate of nonsynonymous changes as are estimated to be in the full proteome, whereas the fraction was considerably below the neutral threshold for somatic and stem cell coding SNVs (Figure 3). A lower fraction of nonsynonymous changes suggests active filtering of SNVs by purifying selection against variants that change the amino acid sequence. Thus, our data suggest that there is no selection happening on germline cells, but there is evidence for purifying selection on somatic and stem cells. If i-cell SNVs are subject to selection, then the selection regime that growing i-cell lines face could select for novel beneficial changes as well as select against deleterious ones ^2,24^. Reef building corals are extremely sensitive to small increases in temperature, but these environmental changes frequently result in the death of just part of a colony. If partial survival of a colony is the result of selection for post-embryonic SNVs in different parts of the colony, then adaptation to environmental change may occur over the lifetime of a single colony. If some of those post-embryonic SNVs are inherited by the surviving polyps’ gametes, then this may be an alternative, rapid route to adaptation for corals.

Our data suggest that anthozoans have i-cells that self-renew and remain multipotent throughout the adult lifespan, which has previously been described in medusozoans. We also show, for the first time, the genome-level consequences of SNVs in i-cells on the mutation load of a long-lived animal species that lacks an embryonic germline. SNVs in the stem cell lines of a coral colony increase the number of SNVs in the sperm by 66%. This may help to explain the high degree of heterozygosity and adaptive polymorphism in many stony coral species. Mechanisms that corals use to avoid mutational meltdown in long-lived cell lineages might include consistent screening by natural selection in proliferating cell lines, or yet-to-be discovered controls on coding gene mutation rates. And, importantly in the context of global climate change, selection in proliferating cell lines may increase the frequency of heritable adaptive polymorphisms in some stony coral species.

## Acknowledgements

EHLN was funded by NSF GRFP and a Stanford Graduate Fellowship in Science and Engineering (Morgridge Family Foundation). Research funded by grants from the Chan-Zuckerberg BioHub and NSF OCE-1736736. We thank N. Neff at CZ-BioHub for sequencing power and A. Voskoboynik P. Bump, and B. Cornwell for stem cell insights. Thanks to R. Ritson-Williams, R. Ross, and BIOTA for field collection support, and thanks to the staff of the Steinhart Aquarium for taking care of the corals. Thanks to S. Matsuda for figure design.

## Methods

### Sample collection

Gravid coral colonies of *Acropora hyacinthus* were collected in Palau (Bureau of Marine Resources permit number RE-19-07 and CITES permit PW19-018) in February 2019 and transported to the Coral Spawning Lab at the California Academy of Sciences where they were kept on a Palauan lunar and day/night cycle until spawning, with methods adapted from ^25^. Colonies were monitored for spawning activity on nights 6 – 9 after the simulated full moon in March 2019 (from 27 March to 30 March 2019). Prior to spawning, pliers were used to break off 2-3 cm branches that were “set,” or showed visual signs of impending gamete release: three branches from each of two colonies, and four branches from a third. Each branch was placed in a labeled 5 mL vial of seawater where they spawned approximately 20 minutes later (Fig. 1). After the gamete bundles were released, they were transferred to labeled 1.5 mL tubes and left to dissociate into eggs and sperm. Upon dissociation, eggs were removed via pipet, leaving a concentrated sperm pool. Each concentrated sperm pool was pipetted into a 1.5 mL tube of RNAlater. Each coral branch was placed in a 5 mL tube of RNAlater. Sperm pools in RNAlater were stored at −20° C and coral branches in RNAlater were stored at −80° C until time for DNA extraction.

### DNA Extraction and Library Preparation

For each coral branch, the top layer of tissue was scraped from the coral skeleton with a razor blade. DNA was then extracted from tissue using the NucleoSpin Tissue Mini kit columns and corresponding protocol for extraction from animal tissue (Macherey-Nagel, Duren, Germany). For each sperm pool, the tube containing RNAlater and sperm was vortexed vigorously, then 200-400 ul of the sperm solution was pipetted out and mixed with 2x volume of D.I. water. The sperm pools were then centrifuged for 3 minutes at 13,000 rpm. The supernatant was pipetted off, leaving just the pelleted sperm at the bottom of the tube. DNA was extracted from sperm pellets using the same Macherey-Nagel NucleoSpin Tissue Mini kit columns and protocol as the parent tissue. Nextera full genome libraries were generated using a modified, low-volume protocol optimized for coral DNA (Supplementary Methods). We constructed two replicate libraries for each DNA extraction (Fig. 1). Libraries were sequenced first on an iSeq 100 for quality control and then on a NovaSeq 6000 S4 at the Chan-Zuckerberg Biohub Sequencing facility in San Francisco, CA, USA.

### Reference genome assembly

In May 2020 we collected sperm from an additional *Acropora hyacinthus* colony for the construction of a high quality *Acropora hyacinthus* reference genome assembly. This colony originated in Palau and spawned at the California Academy of Sciences, where the sperm was collected. Sperm was collected by pipetting, then it was rinsed and spun in seawater 3 times at 13,000 rpm for 3 minutes each spin (following methods from ^26^). The cleaned sperm pellet was then flash frozen in liquid nitrogen. The frozen sperm pellet was shipped to Dovetail Genomics (Scott’s Valley, CA, USA) for DNA extraction, sequencing, and genome assembly. The initial *de novo* assembly was produced through a combination of Illumina short-read sequencing and PacBio long-read sequencing. Proximity ligation was achieved with Dovetail™ Omni-C™ Technology, which uses a sequence-independent endonuclease approach to chromatin fragmentation. The final genome assembly is made up of 908 scaffolds, of which 14 represent full chromosome-length scaffolds, the same number of chromosomes as is in the *Acropora millepora* genome ^27^. The complete assembly is 446,422,234 nucleotides, with N_50_ = 26,527,962 nucleotides.

### Reference genome annotation

Genome annotation was performed using MAKER2^28^ in a *de novo*, iterative approach based on https://gist.github.com/darencard/bb1001ac1532dd4225b030cf0cd61ce2. Transcriptome evidence from *Acropora hyacinthus*^29^ (and https://matzlab.weebly.com/data--code.html), *Acropora millepora*^27,30^, and *Acropora tenuis* (https://matzlab.weebly.com/data--code.html) was provided for the initial round of annotation. Additionally, proteome evidence from *Acropora digitifera*^31^ and *Acropora millepora*^27^ was utilized for the first round. Genome wide repeat families were annotated by RepeatModeler2.0.1^32^ and used as evidence for the initial round. The *ab initio* gene predictors AUGUSTUS v3.2.3^33^ and SNAP^34^ were trained with the gene models annotated by the previous round of annotation. The second round was then conducted with these trained prediction models along with repeat, transcript, and protein evidence annotated during the previous round. A third round of annotation was then performed following the same procedures as round two. Following the final round, the completeness and quality of the annotated transcriptome was assessed with BUSCOv5^35^ and the OrthoDB v10^36^ eukaryota and metazoan datasets. The BUSCO score against the metazoan dataset was 71.3% complete, 13.6% fragmented, and 15.1% missing (Supp. Table 3).

### Read mapping and SNP calling

Adapters were trimmed from reads using Trimmomatic version 0.39. Trimmed reads were mapped to the *Acropora hyacinthus* v1 genome using hisat2 with the parameters --very-sensitive --no-spliced-alignment. Duplicate reads were removed with Picardtools MarkDuplicates. Haplotype calling was performed with the Genome Analysis Toolkit version 4.1.0.0 Haplotypecaller tool^37^. We combined GVCFS from the same coral colony into a multi-sample GVCF using CombineGVCFs. Joint genotype calling was then performed on each mutli-sample GVCF using GenotypeGVCFs with the option –all-sites to produce genotypes for both variant and nonvariants sites^38^. The genotype-called multi-sample VCFs were filtered with SelectVariants to filter files by depth, with minimum depth and maximum depth determined by a Poisson distribution of the average depth for a given sample, with p <0.0001^39^. The filtered files resulting from these steps were considered the “callable” regions of the genome, and were used as the denominator for mutation frequency calculations. We filtered for just biallelic single nucleotide polymorphisms (SNPs) using VCFtools. For the complete read mapping and SNP calling pipeline see https://github.com/eloralopez/CoralGermline

### Identifying post-embryonic single nucleotide variants (SNVs)

Single nucleotide variants from the genotyped colony VCFs using custom Python3 and R scripts (https://github.com/eloralopez/CoralGermline). Putative post-embryonic SNVs were identified by comparing the parent branch genotype calls from a given colony. A SNP was called a putative post-embryonic SNV if the SNP a.) appeared in just one branch of the colony, and b.) the SNP had the same genotype call in both replicate libraries from that mutant branch (Fig 1c, d).

Germline mutations were identified by comparing the sperm genotype calls from a given colony. A SNP was called a putative unique germline mutation if the SNP a.) appeared in just one sperm pool spawned by the colony, b.) the SNP had the same genotype call in both replicate libraries from the sperm pool, and c.) the genotype in the mutant sperm pool did not match the genotype of the parent branch that spawned it (Fig. 1). Alternatively, we called a SNP a putative global germline mutation if the SNP a.) appeared in every replicate library from every sperm pool spawned by the colony and b.) the genotype in the sperm pools did not match the genotypes of any of the parent branches in that colony (Fig. 1).

### Classifying putative SNVs

Once we had generated a set of putative somatic and germline mutations, we classified each SNV as either a Gain of Heterozygosity (GoH) or Loss of Heterozygosity (LoH) mutation, and classified the directionality of the change (A to T, etc.) as described in ^14^.

### Final filtering of putative mutations to arrive at final set of mutations

The final filtering step was to eliminate putative mutations that had been classified GoH if the putatively mutant allele had been seen before in any of the other libraries. This step was necessary because the filtering threshold to call a heterozygotes was 10%. This means that if the putatively mutant allele was present at <10% allele frequency in other samples, it was not truly a mutant, but present in the colony at low levels. Similar, we eliminated putative mutations classified as LoH if the mutant sample was heterozygous at a level <10%. The mutations that passed this filter is the final set of mutations upon which all subsequent analyses were performed.

### Are parent SNVs shared by their sperm pool or not?

Once we arrived at a set of somatic mutation with all filters applied, we checked to see if the genotype of the mutant parent matched the genotype of the sperm pool that came from that branch. GoH SNVs were considered SHARED if one or both genotypes of the two sperm pool replicates spawned by the mutant branch matched the mutant genotype and NOT SHARED if neither of the genotypes of the two sperm pool replicates spawned by the mutant branch matched the mutant genotype (Fig. 1). LoH SNVs were considered SHARED if both genotypes of the two sperm pool replicates spawned by the mutant branch matched the mutant genotype and NOT SHARED if one or neither of the genotypes of the two sperm pool replicates spawned by the mutant branch matched the mutant genotype (Fig. 1).

### Designating SNV effects on codons

We classified each mutation by the genomic region (intron, exon, etc.) it fell in and, if it fell in a coding region, the type (synonymous, missense) using the program snpEff^40^ configured with the *Acropora hyacinthus* version 1 genome. We calculated the rate of missense and synonymous SNVs per bp in the coding region by dividing the number of SNVs by the total callable coding region. We calculated the average percent of coding SNVs that were missense per sample for three categories: Parent Only, Parent and Sperm, and Single Sperm Pool Only. There were no coding SNVs in the All Sperm Pools category (Extended Data Fig. 6).

### Selection on mutations

To determine whether the percent of missense coding SNVs in the Parent Only, Parent and Sperm, and Single Sperm Pool Only categories were significantly different from what would be expected under neutrality, we estimated the percent missense sites across the full *Acropora hyacinthus* proteome using a custom Python script that counts how many of each type of amino acid are present in the full proteome (https://github.com/eloralopez/CoralGermline).

### Mutation rates

To catalogue post-embryonic SNVs, we compared sequences from 3 parent branches in each colony and recorded cases in which one branch showed a genotype different from all the others. This catalogue includes true somatic mutations in that branch, but also includes any sequencing or PCR errors injected during sample preparation and sequencing. To limit these errors, we compared sequences from each technical replicate and catalogued a mutation only when it was visible in both replicates of a branch and in none of the replicates of any other branch. Comparing technical replicates was highly successful at reducing noise from sequencing error, eliminating over 90% of putative SNVs that would have been identified using the same pipeline and filters without technical replicates (Extended Data Figs. 2, 3). To find the average frequency per nucleotide of somatic mutations unique to a given branch in a coral colony, we divided the number of SNVs in a sample by the number of total callable nucleotides sequenced for that sample (Supplementary Table 1).

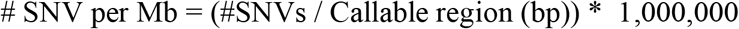

The callable genome sizes were 1.2 x 10^8^ bp, 1.2 x 10^8^ bp, and 0.80 x 10^8^ bp and the callable coding region sizes were 1.3 x 10^7^ bp, 1.3 x 10^7^ bp, and 0.91 x 10^7^ for each of the three colonies, CA56, CA60, and CA65, respectively (Supp. Table 1).

### Tissue mixtures

Because DNA was extracted from tissue scrapings encompassing multiple polyps per parent branch, we were initially concerned that the parent sample might be a heterogenous mix of somatic and germ tissues. Germ and stem cells in other cnidarians tend to reside at the base of the polyp, so the scraping method was intended to take off just the somatic tissue and not the germ or stem cells. To check this, we plotted the average variant allele frequency (VAF) of the mutant parent (that is, the average VAF for the two technical replicate libraries) against the average VAF of the mutant sperm pool.

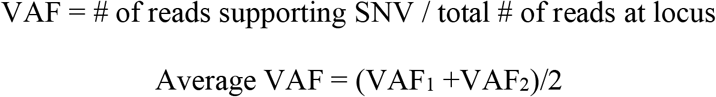

If the parent samples were a mix of somatic and germ and/or stem cells, then a germline mutation may have erroneously been called a somatic mutation. If that were the case, then the VAF of the parent would be considerably lower than the VAF of the sperm pool, because the sperm pool’s mutant would not be diluted by nonmutant somatic tissue. In that case, we would expect the trendline of the parent VAF: sperm VAF linear model to have a slope significantly greater than 1. In reality, the slope of the trendline was less than 1, suggesting that we undercounted the number of mutant reads in the sperm. This gives us confidence that the parent samples were in fact all or almost all somatic tissue, and that the mutations found in the parents were in the soma. We used a linear regression model to check the relationship between the variant allele frequency of the mutant parent and the variant allele frequency of the mutant sperm for all inherited GoH mutations, where variant allele frequency equals the number of reads with mutation divided by the total number of reads. The slope of the trendline was 0.36, with R = 0.3 and p = 5.9e-08 (Extended Data Fig. 7). This suggests that there may have been some undercounting of the number of mutant reads in the sperm. The distribution of average VAF in the parents for SNVs classified as “Parent Only” has a larger leftward skew, which also suggests that some SNVs found at low frequency in the parent samples may have been missed in the sperm (Extended Data Fig. 8). This gives us confidence that the parent samples were in fact all or almost all somatic tissue, and that the mutations found in the parents were in the soma.

### Types of mutations

For Gain of Heterozygote mutations (where the mutant genotype is a novel heterozygous SNP) we classified the somatic mutation as inherited by the sperm if at least one of the technical replicate libraries of the sperm contained the mutant allele. If neither sperm library contained the mutant allele, then it was considered not inherited by the sperm. For Loss of Heterozygosity mutations (where the mutant is a homozygous genotype at a site for which the parent colony is heterozygous) to be classified as inherited by the sperm both sperm libraries had to be entirely homozygous for the allele of the mutant parent. For these cases, if any read for a parent sample showed the minor allele also seen in the putative GoH mutation, we did not call these as mutations. Likewise, if any of the sperm reads showed the minor allele also seen in the parent branch, we did not call this a LoH mutation. These stringent filters reduced the number of mutations we could characterize but led to high confidence in our characterization of variants.

## Data availability

All raw fastq files, as well as the *Acropora hyacinthus* genome version 1 assembly, are accessioned under BioProject PRJNA707502 at NCBI. The accession numbers for the fastqs are SAMN18207983-SAMN18208014 and the accession number for the assembly FASTA is SAMN20335437.

## Code availability

The code used for this study can be found at **https://github.com/eloralopez/CoralGermline.**

## Author contributions

EHLN, RA, and SRP designed the study. RA kept corals alive and healthy and facilitated lab spawning. EHLN and RA collected parent and sperm samples for this study. EAS optimized the library preparation protocol used in this study. EHLN extracted DNA and constructed libraries for all samples. EHLN designed and performed all bioinformatic analyses for the study, except for genome annotation, which was performed by EAH. EHLN wrote the manuscript and all authors provided input and feedback on drafts.

## Additional Information

Supplementary information is available for this paper:

Supplementary Table 1. Summary of sequencing depth and SNV counts (.txt file)
Supplementary Table 2. List of verified SNVs and their metadata (.txt file)
Supplementary Table 3. Summary statistics for *Acropora hyacinthus* version 1 genome annotations (.xlsx file)
Supplementary Methods. The optimized full genome library preparation protocol for corals (.docx file).

## Extended data figure/table legends

**Extended Data Figure 1.**
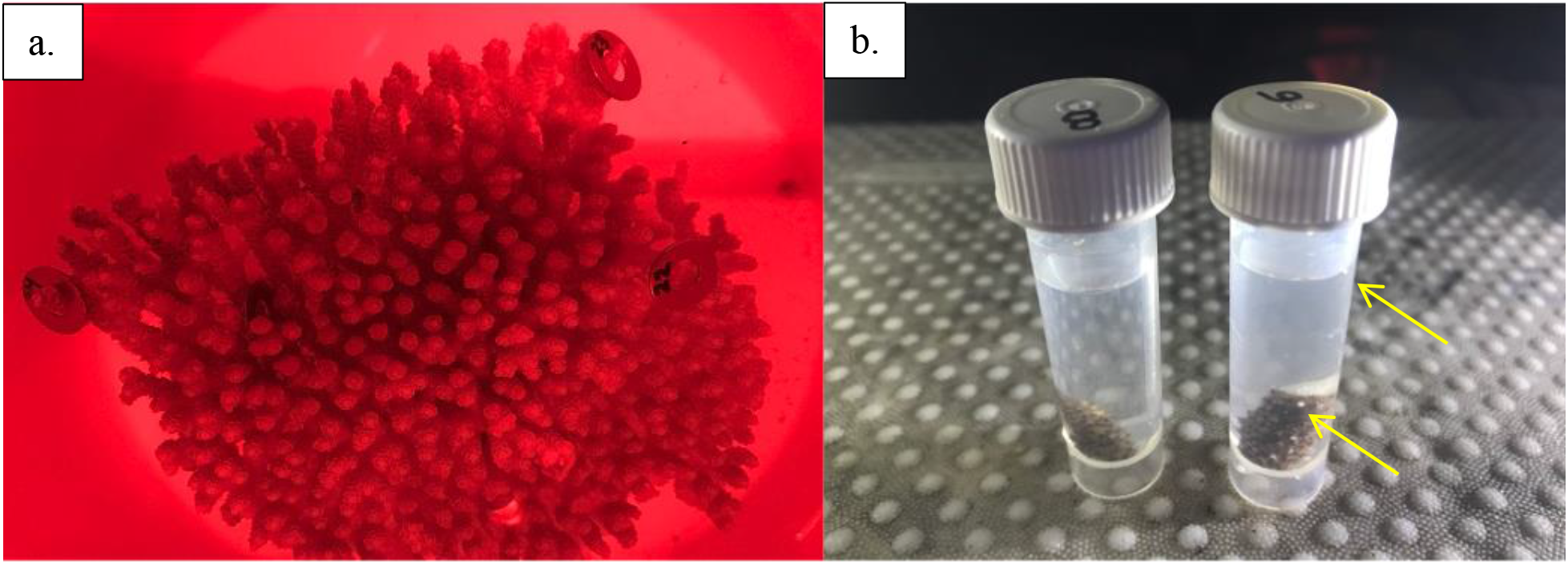
Photos of the experimental setup. a.) Colony 60, with sampled branches denoted by numbered washers. For scale, the outside diameter of each washer is 1.59 cm. b.) Two of the branches (CAP6 and CAP8) sampled from Colony 56, releasing gamete bundles (indicated by arrows) into their respective 5 ml tubes of seawater.

**Extended Data Figure 2.**
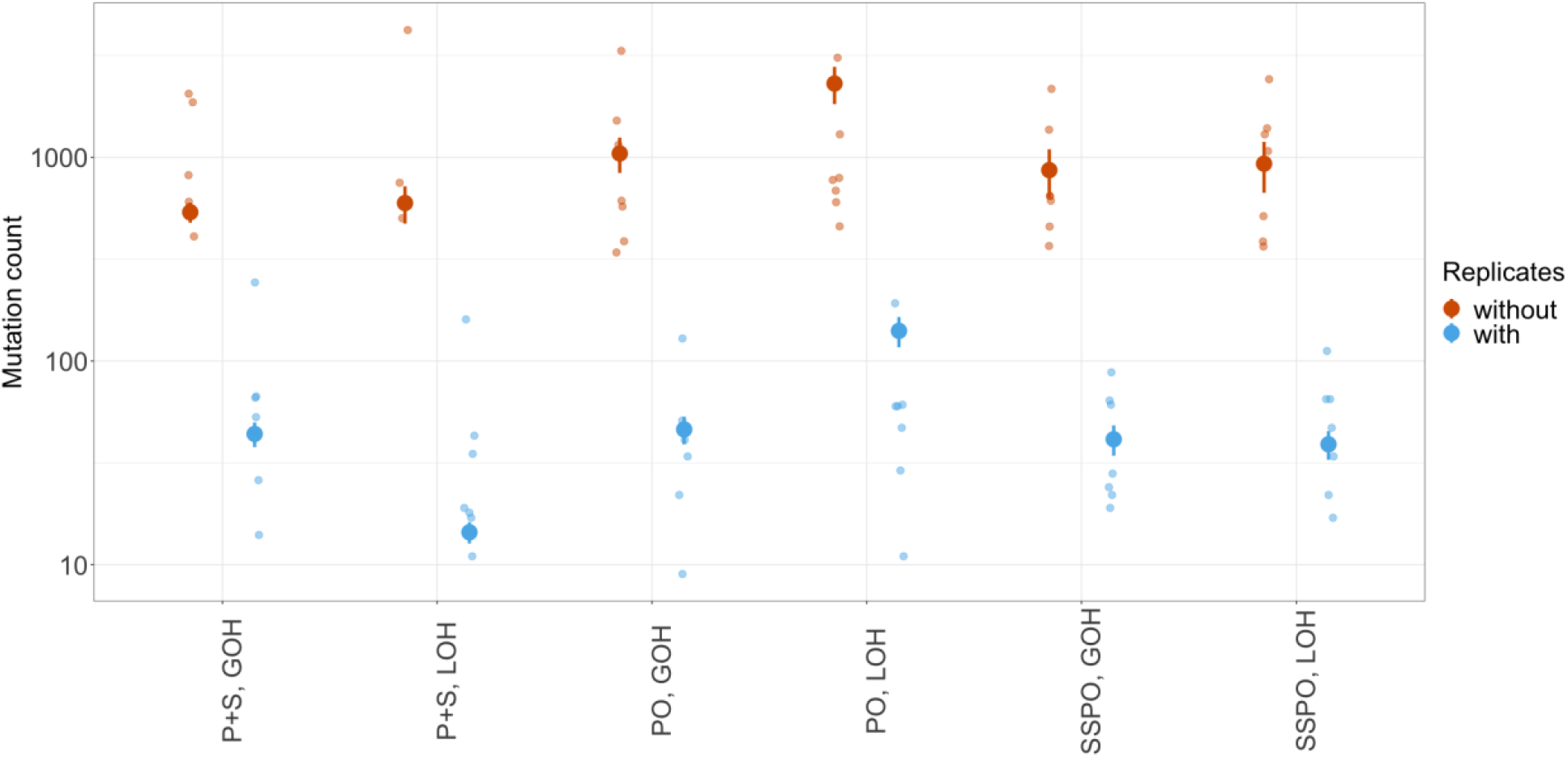
The mean number of mutations identified in each parent branch in the Parent Only (PO) or Parent and Sperm (PO), or single sperm pool only (SSPO) when looking at one library per sample (aka without technical replicates, pink) or two libraries per sample (with technical replicates, gray). Error bars represent ± 1 s.e.m. Each data point (N=7) is shown as a smaller point for each category. Note that the y axis is on a log-10 scale.

**Extended Data Figure 3.**
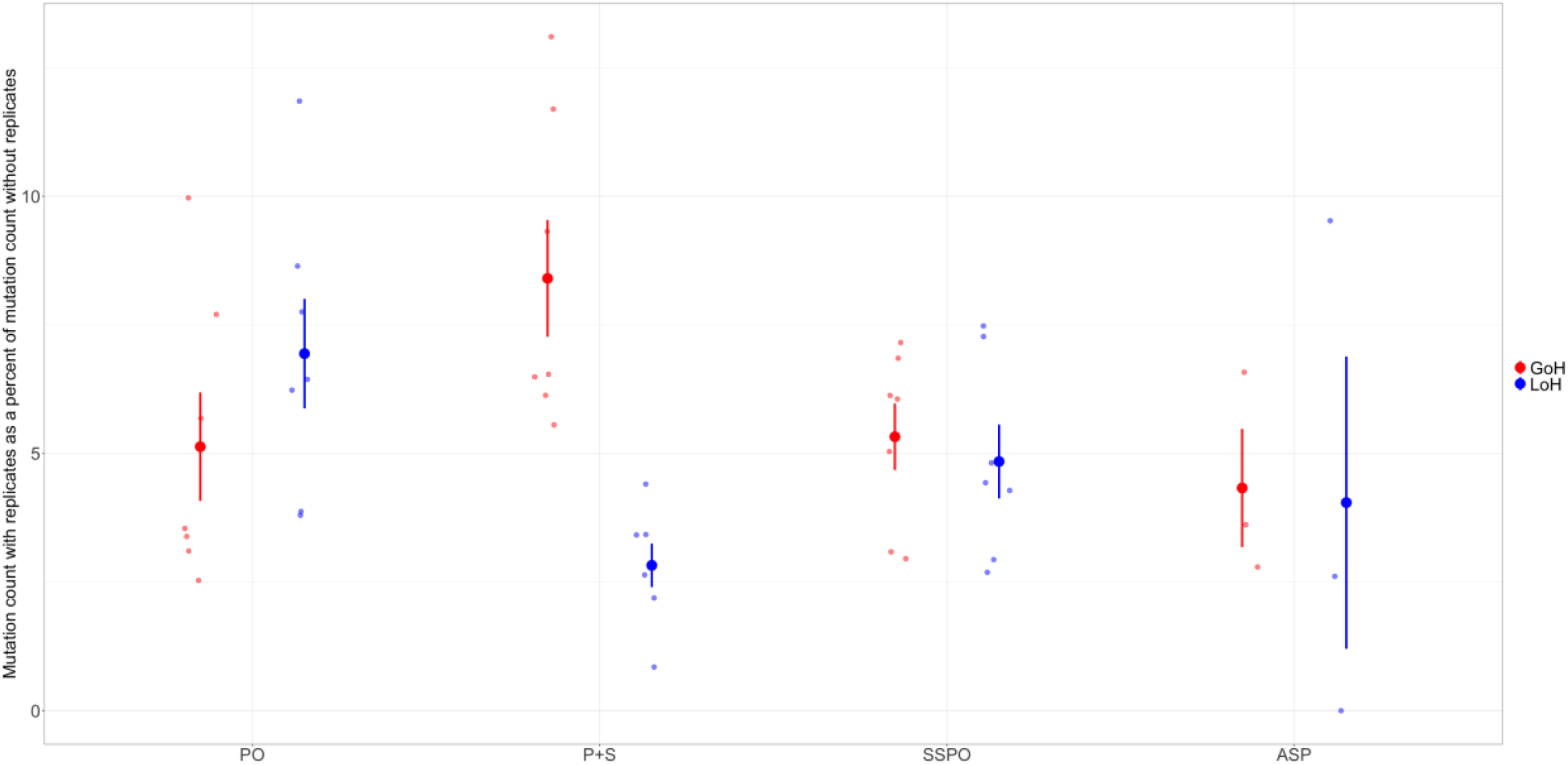
The percentage of mutations identified when replicate libraries are included for each of the four categories of SNVs: Parent Only (PO), Parent and Sperm (P+S), Single Sperm Pool Only (SSPO), and All Sperm Pools (ASP). Error bars represent ± 1 s.e.m. Each data point (N=7) is shown as a smaller point for each category. The percentage is expressed as a fraction of the number of mutations identified when no technical replicates are included, such that: % = (# of mutations identified using technical replicates*100) / (# of mutations identified without technical replicates)

**Extended Data Figure 4.**
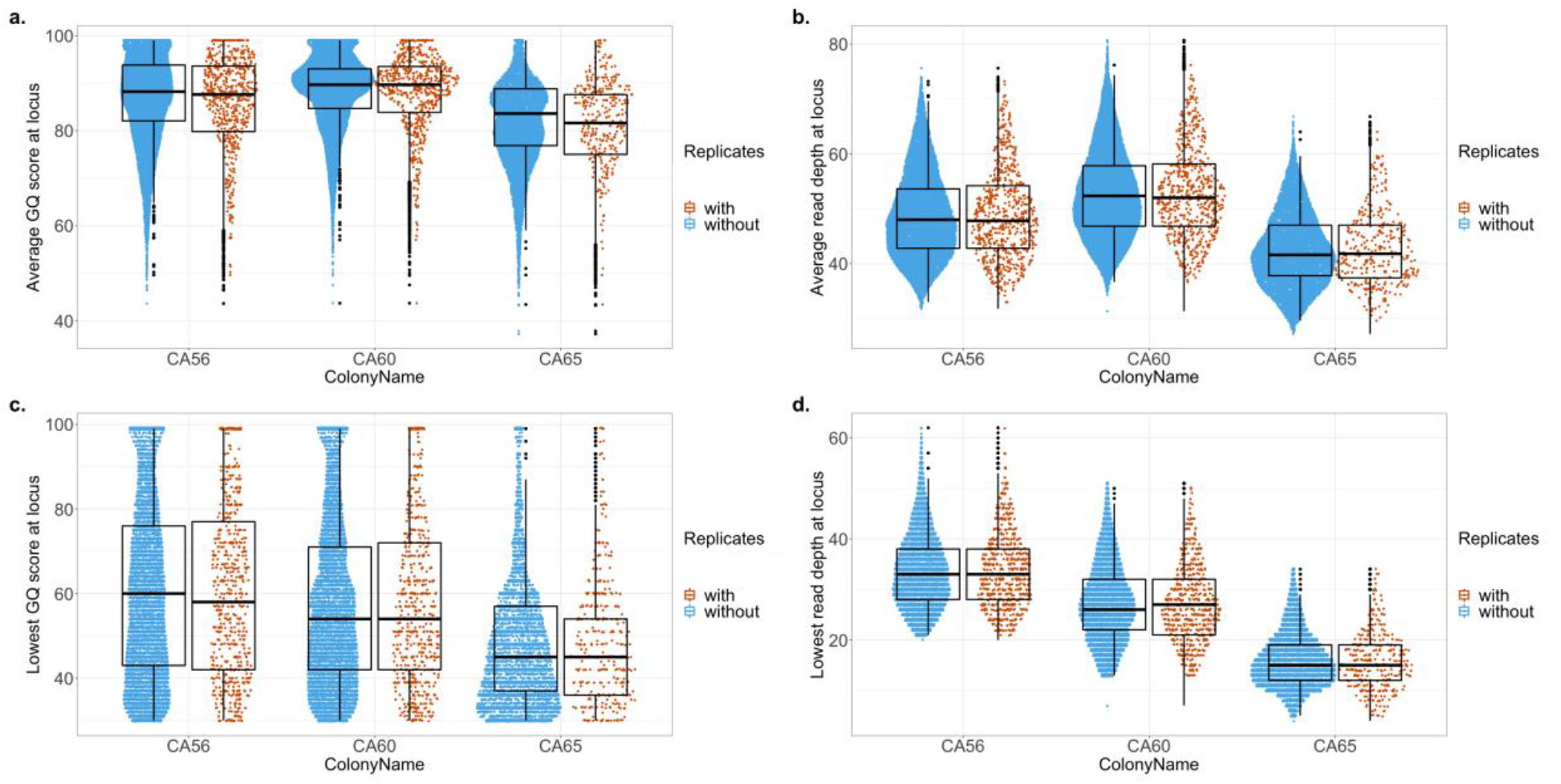
Distribution of average GQ score and average read depth for each mutation in the colony (top) as well as distributions of the lowest GQ score and lowest read depth for every mutation identified in each colony (bottom). Mutations found when using no technical replicates are shown in pink, and the mutations found when including technical replicates shown in gray.

**Extended Data Figure 5.**
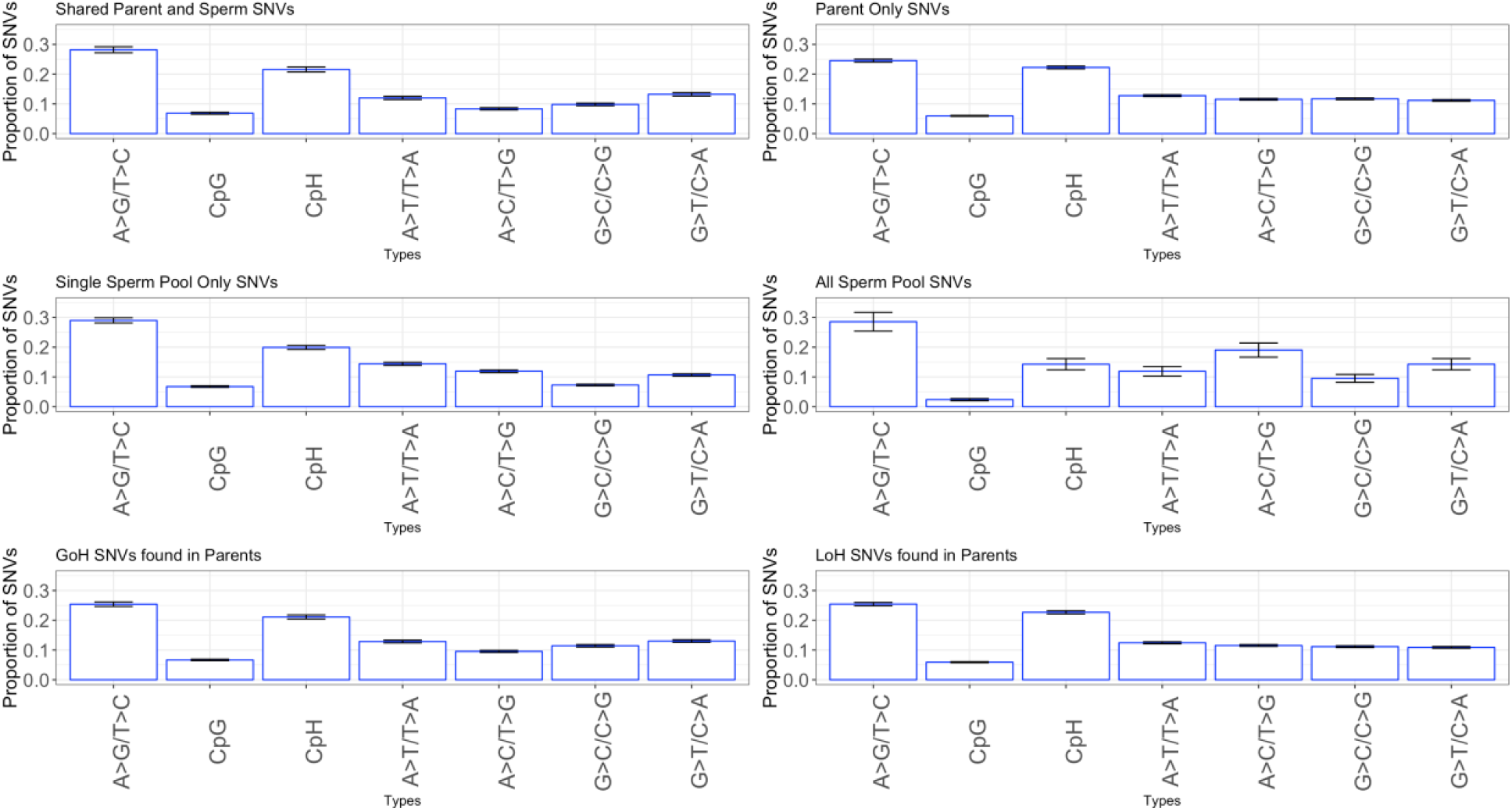
SNV spectra from various subsets of SNVs. None of the SNV spectra were significantly different each other for the different data subsets (see *X*^2^ tests below). Error bars represent ± 1 s.e.m. *X^2^* tests of independence results for Extended Data Figure 5:

| Comparison | $X^2$ |
| --- | --- |
| Parent only vs. Shared Parent and Sperm | df = 6, N=1715, $X^2 = 7.3495$, p= 0.2897 |
| Single Sperm Pool Only vs. All Sperm Pools | df = 6, N=604, $X^2 = 4.2981$, p= 0.6364 |
| Parent GOH SNVs vs Parent LOH SNVs | df = 6, N=1715, $X^2 = 3.9411$ p= 0.6846 |

**Extended Data Figure 6.**
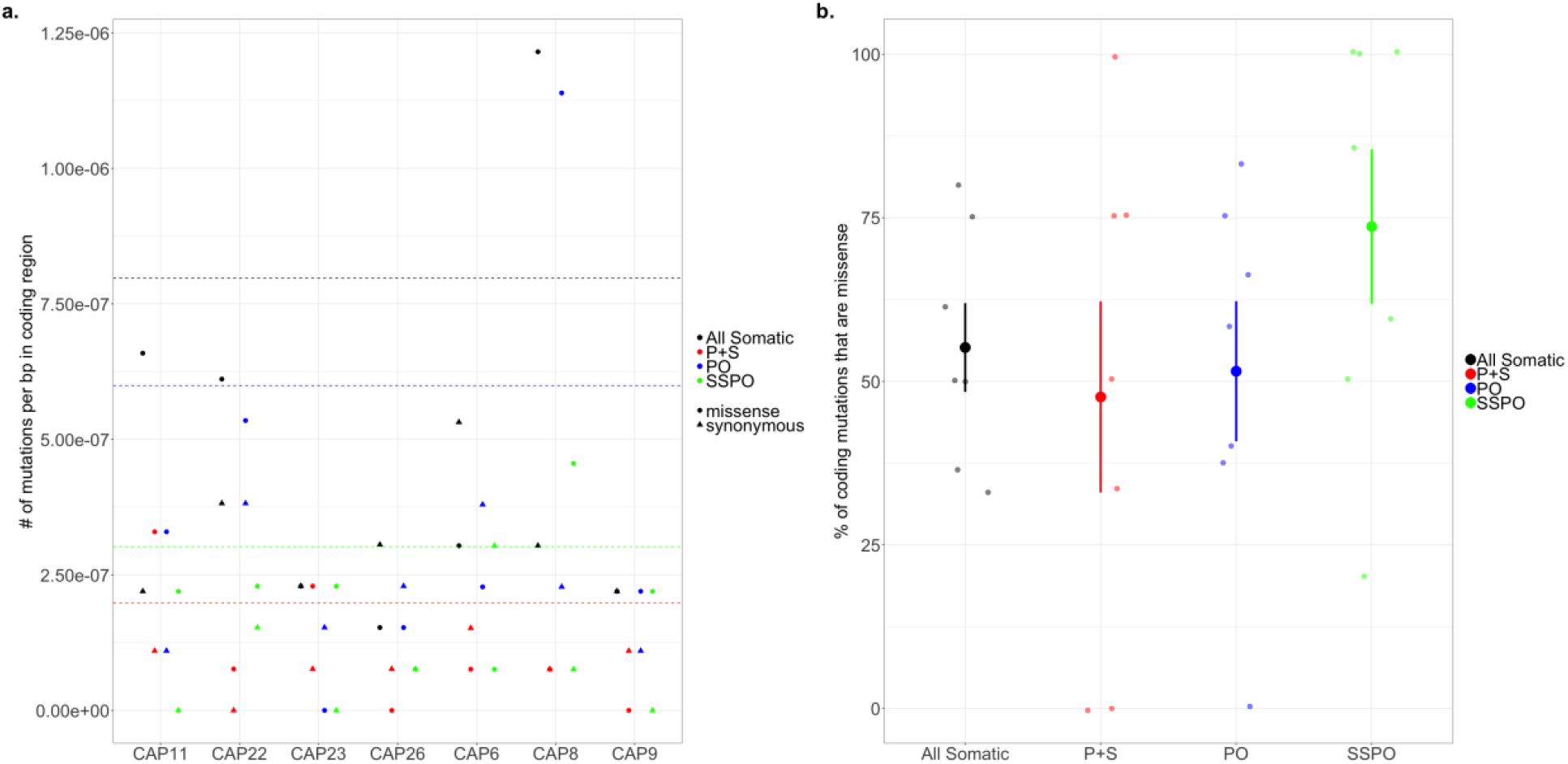
The number of missense (circle) and synonymous (triangle) SNVs per bp of the coding region for each category (all somatic, P+S, PO, and SSPO) for each sample. Dashed lines indicate the mean number of coding SNVs (the sum of missense and synonymous) for each category across all samples.

**Extended Data Figure 7.**
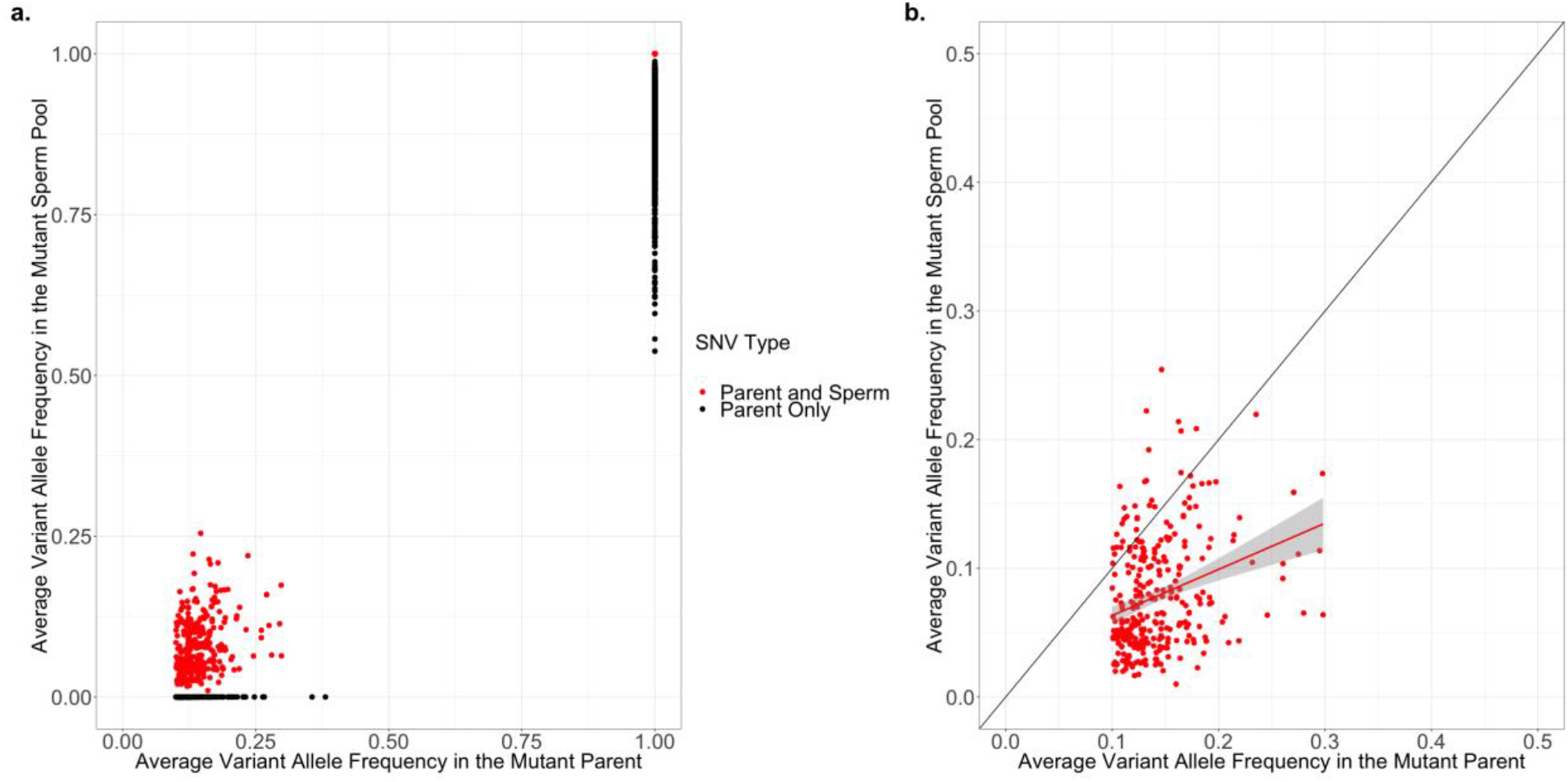
Variant allele frequencies for parents and their respective sperm pools. a.) GOH mutations for which sperm VAF =0 were classified as Parent Only (black), likewise LOH variants in the parent for which the VAF < 1 in the sperm were classified as Parent Only. All other SNVs were classified as being shared by both a parent branch and its corresponding sperm pool (red). B.) For GOH SNVs that are shared by a parent branch and its sperm, average variant allele frequency of the two replicate parent libraries is positively correlated with the average variant allele frequency of the two replicate sperm pool libraries. The slope of the relationship is 0.36 and R=0.3 (red line). The black line shows a 1:1 line for reference.

**Extended Data Figure 8.**
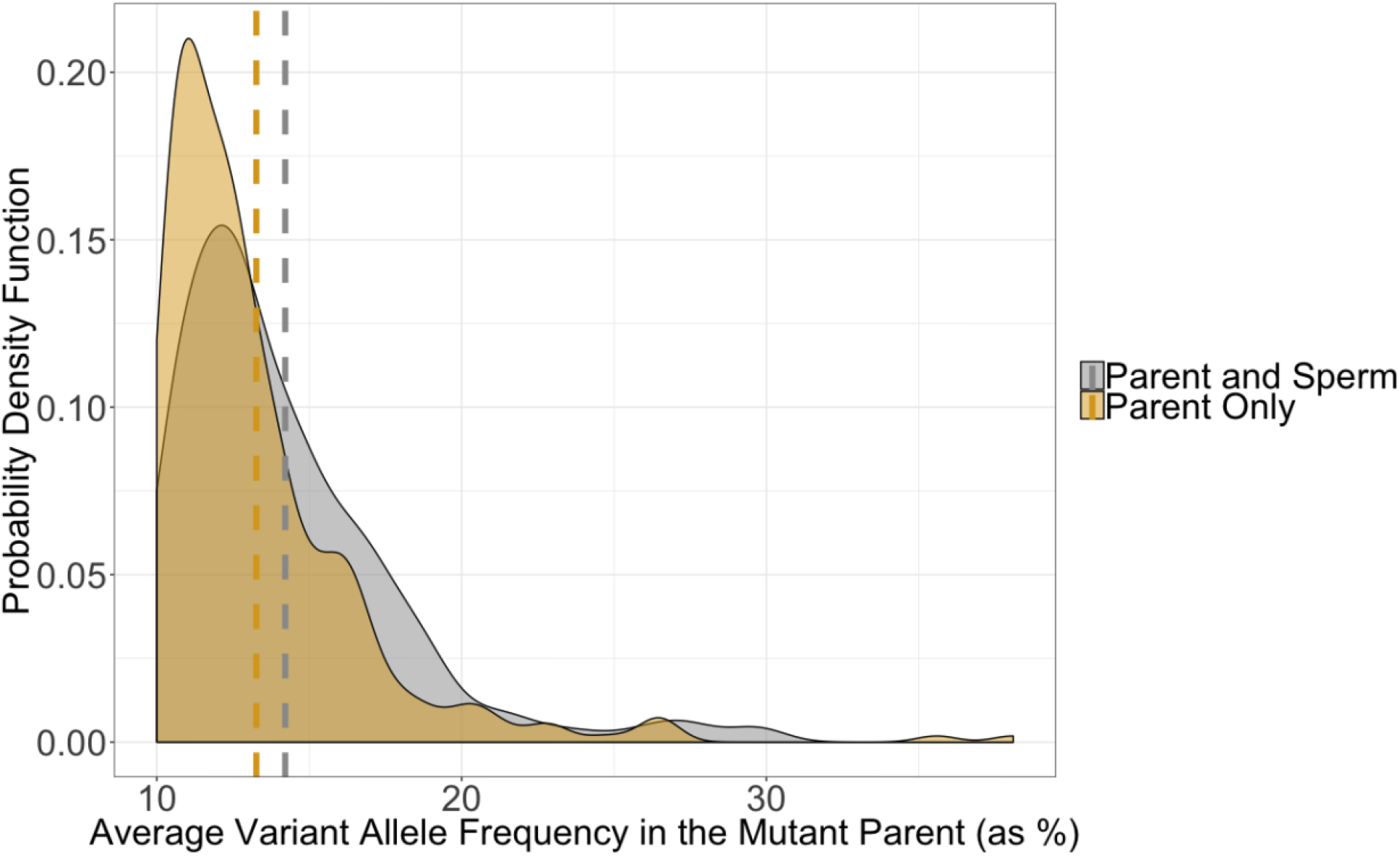
The distribution of variant allele frequencies in the parent branches for GOH SNVs in the parents only (orange) and in the parent and sperm samples (gray). The mean parent variant allele frequency for each category is marked by dashed lines.

**Extended Data Table 1.** Comparison of SNV frequencies in the coding region vs. across the full genome for six SNV types (shown in Figure 2c). Error shown is ±1 s.e.m.

| Type | Average frequency per bp<br>(Full genome, coding) | Wilcox test (two-tailed,<br>paired) |
| --- | --- | --- |
| Parent Only, GOH | Full: $4.4 \times 10^{-7} \pm 0.6 \times 10^{-7}$<br>Coding: $9.2 \times 10^{-8} \pm 3.1 \times 10^{-8}$ | V=28, P= 0.016 |
| Parent Only, LOH | Full: $1.32 \times 10^{-6} \pm 0.19 \times 10^{-6}$<br>Coding: $5.2 \times 10^{-7} \pm 1.4 \times 10^{-7}$ | V=28, P= 0.016 |
| Parent and Sperm, GOH | Full: $4.8 \times 10^{-7} \pm 0.9 \times 10^{-7}$<br>Coding: $1.8 \times 10^{-7} \pm 0.6 \times 10^{-7}$ | V=27, P= 0.031 |
| Parent and Sperm, LOH | Full: $2.6 \times 10^{-7} \pm 1.1 \times 10^{-7}$<br>Coding: $1.6 \times 10^{-8} \pm 1.6 \times 10^{-8}$ | V=28, P= 0.016 |
| Single Sperm Pool Only,<br>GOH | Full: $3.8 \times 10^{-7} \pm 0.5 \times 10^{-7}$<br>Coding: $1.5 \times 10^{-7} \pm 0.5 \times 10^{-7}$ | V=28, P= 0.016 |
| Single Sperm Pool Only,<br>LOH | Full: $3.6 \times 10^{-7} \pm 0.4 \times 10^{-7}$<br>Coding: $1.6 \times 10^{-7} \pm 0.6 \times 10^{-7}$ | V=27, P= 0.031 |

