## Supplementary Methods for "Mutations in coral soma and sperm imply lifelong stem cell renewal and cell lineage selection"

**Supplementary Methods for López-Nandam et al.**

Illumina Sequencing Library Preparation

Protocol

Modified from the Illumina’s Nextera Protocol by Sergey Kryazhimskiy (Desai Lab)

Updated by Beth Sheets for the Palumbi Lab (last update: November 27, 2018 )

**Helpful background literature:**

- Therkildsen & Palumbi (2015) Practical low-coverage genome-wide sequencing of hundreds of individually barcoded samples for population and evolutionary genomics in non-model species
- Baym et al. (2014) Inexpensive Multiplexed Library Preparation for Megabase-Sized Genomes

**Consumables**	**Product Supplier Cat. #**

Nexterta TDE1 Tagment DNA enzyme	Illumina 15027865 (120uL)

Nexterta Tagment DNA buffer	Illumina 15027866 (600uL)

Nextera Index Kit (96 indices, 384 samples)	Illumina FC-121-1012

Library Amplification Kit KAPA Biosystems	KK2611/KK2612

AMPure XP beads Beckman Coulter	A63880

Foil Seal for recovering Illumina index plate

Primer P1 (AATGATACGGCGACCACCGA), purified

with HPSF, at 10μM

Primer P2 (CAAGCAGAAGACGGCATACGA), purified

with HPSF, at 10μM

Labware

• sterile 8-strip tubes & caps

• 200μl and 10μl multichannel pipettes

• 96-well plate magnetic stand (e.g., Life Technologies, Cat. #123-31D)

• freshly prepared 80% ethanol (e.g. within 2 days)

• sterile HPLC water

Best practices

- I find doing each reaction in an 8-strip tube is easiest to view and manipulate beads on the magnet
- Use filter tips before first PCR to reduce contamination
- Make sure Ampure beads are at room temperature before use & thoroughly mix them by vortexing
- When mixing sample with beads, pick up liquid from bottom and drop near top of liquid until coloration is homogenous. Cap and periodically mix gently during 2min incubation
- When separating small volumes from beads in a plate with circular magnets, I gently tilt 8-strip tube towards back of circle magnet and wait for beads to accumulate on the back side of the tube. I then slide the tip of my pipet down the front side of the tube.
- Pick out indices before starting. For 1X coverage genomes, we usually run 60 samples per lane. Confirm you have enough index primers.

1. Input DNA

The input DNA for this protocol needs to be very precise because the tagmentation enzyme is quite sensitive to DNA concentration and will yield different size product smears with different input concentrations. We input 2ng total gDNA in 2uL of solution, so samples should be diluted precisely to 1ng/uL. Here are my recommendations for easier quantification:

1. After DNA extraction, elute samples into 10mM Tris-HCL buffer with EDTA so that our stock samples do not degrade. We dilute the samples going forward so that the EDTA should not affect our downstream processes.
2. After extraction, before Qubit, dilute stock samples into working solution with 10uL of gDNA into 100uL of HPLC water so you are able to Qubit with higher volumes of input DNA to reduce pipetting error.
3. Qubit working solution with 3uL of input DNA to reduce pipetting error.
4. Using the working solution, run DNA on an agarose gel (1.5%, 8uL DNA 4uL Gel Red) to check the quality of the DNA – only high molecular weight DNA will work! If the sample is degraded, re-extract and check its quality again before proceeding.
5. For high molecular weight samples, use C1V1=C2V2 to calculate a dilution that yields exactly 1ng/uL concentration (try to keep the pipetting range above 5uL to reduce pipetting error). Check that this dilution worked! If not, do a second serial dilution and check again.
2. Tagmentation of genomic DNA

Background

The Illumina tagmentation enzyme (TBE1) randomly fragments the genome and “tags” the cleaved DNA ends with a universal overhang that match a segment of the P1 and P2 primers in this protocol. The longer the tagmentation reaction goes at 55C, the more DNA is cleaved and tagged. We use a 1/10 reaction volume from the original protocol because this seems more stable.

Preparation

1. Prepare gDNA at a concentration at 1 ng/μl

2. Confirm the concentration by HS Qubit assay

3. Remove the TD, TDE1 and gDNA from the –20°C and thaw on ice

4. After thawing, mix all reagents and gDNA by gently vortexing

Procedure (samples (n) = rows (r) x columns (c) )

1. Heat thermocycler up on a paused 55C in the “Tagmentation” protocol.
2. Make the Tagmentation Master Mix (TMM) by mixing n x 1.02 x 2.5 μl of TD Buffer and n x 1.02 x 0.5μl of TDE1 in a PCR tube. Mix thoroughly by gently pipetting the mixture up and down 20 times
3. Distribute TMM into r tubes (or a PCR strip), c x 1.02 x 3μl into each tube
4. With a multichannel pipette, distribute TMM into all wells of a fresh plate (“tagmentation plate”), 3μl per well
5. With a multichannel pipette, transfer 2μl of gDNA into the tagmentation plate (total volume = 5 μl per well). Mix by gently pipetting up and down 10 times. Change tips after every transfer.
6. Cap the tubes
7. Give the plate a quick spin to collect all liquid at the bottom (~1000 rpm for 1 min). Do not forget to balance the centrifuge.
8. Place the plate in the thermocycler and run the following program:

• 55°C for 3 min

• Hold at 10°C

NOTE: Ensure that the lid is tight and that it is heated during incubation

3. PCR (with reconditioning)

Background

This step adds your i5 and i7 sample identification indices to the overhangs of your DNA fragments. For ~60 samples/lane you can add any combination of unique indices because this number will generate plenty of diversity on the flow cell. For less than 12 samples/lane, consult the Illumina index guide.

Preparation

1. Remove the KAPA polymerase mix (KAPA amplification kit KK2611/KK2612) and the 96 well plate of indices from the –20°C and thaw on ice.
2. After thawing, mix reagents and indices by vortexing.
3. Bring AMPure XP beads to room temperature
4. Prepare fresh (don’t use old) 80% ethanol from absolute ethanol in a sterile reservoir (50mL of 80% ethanol is 42ml of 95% ethanol to 8ml HPLC H20).

Procedure (samples (n) = rows (r) x columns (c))

1. Add 7.5μl of 2x KAPA to each well of the tube with the tagmentation reaction. Mix thoroughly by gently pipetting the mixture up and down 20 times.
   a. You can also aliquot the KAPA into a strip tube by placing 7.5 x 1.02 x no. of strip tubes of KAPA into each well of the strip tube and then use a multi-channel to add 7.5 μl KAPA to each well.
2. Transfer 2.5 μl of the already combined N5xx/N7xx index into its corresponding well (you need to decide which indices to label each sample with according to what is being combined on a sequencing lane). Mix thoroughly by pipetting up and down.
3. Cap each tube. Make sure to press well on each well.
4. Give the plate a quick spin to collect all liquid at the bottom at 1000 rpm for 1 min.
5. Place the tubes in the thermocycler and run the following program:

72°C for 3 min

98°C for 2:45 min

98°C for 15 sec

62°C for 30 sec

72°C for 3 min

Repeat steps (3–5) 8 times

72°C for 1 min

Hold at 4°C

NOTE: Ensure that the lid is tight and that it is heated during incubation

6. After PCR completes, remove primer dimer using a 1X Ampure bead ratio clean-up.
7. Check that the volume of your PCR product is at 15uL. Increase volume to 30μl by adding 15 μl (or adjust if necessary to get to 30).
8. Add 30 μl of room temperature, well-mixed Ampure beads to each well containing 30 μl of PCR product. Mix well by pipetting up and down 20 times and liquid is a homogenous color.
9. Let mixture incubate for 5 minutes (not on the magnetic stand). Lightly spin tubes down to remove any product on the sides.
10. Place the strip tubes on the magnetic stand and incubate for 1 min to separate bands from solution. Wait for the solution to become clear.
11. While the tube is on the magnetic stand, aspirate clear solution from the tube and discard (~60uL). Do not disturb the beads. If beads are accidentally pipetted, resuspend them back, wait for solution to clear, and repeat.
12. While tubes are on the magnetic stand, dispense 200 μl of freshly prepared 80% ethanol into each well and incubate for 30 seconds at room temperature. Aspirate out ethanol without disturbing the beads and discard. Repeat for a total of 2 washes.
13. Remove any remaining ethanol with a P10 pipette.
14. Let the tubes air dry for approximately 5 min. Do not over dry beads.
15. Take the tubes off the magnetic stand. Add 16 μl of HPLC water to each well by dropping the liquid onto the beads dried to the side of the tube. Carefully resuspend the beads by mixing 10-15 times. Incubate for 5 min at room temperature. DNA is now in solution.
16. Place the tubes back onto the magnetic stand and incubate for 1 min to separate beads from solution. Wait for the solution to become clear.
17. While the tube is on the magnetic stand, aspirate 15 μl of clear solution from the tubes and transfer to a fresh tube. Do not disturb the beads. If beads are accidentally pipetted, resuspend them back, wait for the solution to clear up, and repeat.
18. Make Reconditioning PCR Master Mix (RMM), by mixing n x 1.02 x 17μl of KAPA polymerase mix, n x 1.02 x 1μl of primer P1, and n x 1.02 x 1μl of primer P2. Mix thoroughly by gently pipetting the mixture up and down 20 times.
19. Distribute RMM into a PCR strip tube, c x 1.02 x 19μl into each tube
20. With a multichannel pipette, transfer 19 μl of RMM into each well of the plate (final PCR volume 36μl). Mix by gently pipetting up and down 10 times. Change tips after every transfer.
21. Cap each tube and make sure they fit well.
22. Give the tubes a quick spin to collect all liquid at the bottom at 1000 rpm for 1 min.
23. Place the tubes in the thermocycler and run the following program:

95°C for 5 min

98°C for 20 sec

62°C for 20 sec

72°C for 2 min

Repeat steps (2–4) 4 times

72°C for 2 min

Hold at 4°C

NOTE: Ensure that the lid is tight and that it is heated during incubation

4. PCR Clean-up and size selection

Preparation

1. Bring AMPure XP beads to room temperature

2. Prepare fresh 80% ethanol from absolute ethanol in a sterile reservoir.

Procedure (n samples = r rows, c columns)

1. Centrifuge the plate to collect all liquid (1000 rpm for 1 min)

2. Vortex beads for 30 sec to ensure that they are evenly dispersed

3. Check the volume of the PCR solution -- it should be at 34 μl, but PCR evaporation may reduce the volume. If it is not at 34 add enough HPLC water to get the volume to 34 μl (this is important to get the correct 0.65 ratio of Ampure beads).

3. Transfer c x 1.05 x 1 x 22.1 μl of Ampure beads into a a PCR strip

4. Using a multichannel pipette, transfer 22.1 μl of beads into each well containing the PCR

product (+water). Mix well by gently pipetting up and down 20 times. The color of the mixture

should appear homogeneous after mixing. Change tips between columns

5. Incubate at room temperature for 5 min. DNA is now on the beads

6. Place the plate on the magnetic stand and incubate for about 1 min to separate beads from

solution. Wait for the solution to become clear

7. While the plate is on the magnetic stand, aspirate clear solution from the plate and discard.

Do not disturb the beads. If beads are accidentally pipetted, resuspend them back, wait for

the solution to clear up, and repeat

8. While the plate is on the magnetic stand, dispense 200μl of 80% ethanol into each well and

incubate for 30 seconds at room temperature. Aspirate out ethanol without disturbing the

beads and discard. Repeat for a total of 2 washes

9. Remove any remaining ethanol with P10 pipette.

10. Let the plate air dry for approximately 5 min. Do not overdry the beads.

11. Take the plate off the magnetic stand. Add 21 μl of HPLC or 10mM Tris-HCl (pH 8) to each well of the plate. Carefully resuspend the beads by mixing 10-15 times. Incubate for 5 min at room

temperature. DNA is now in the solution

12. Place the plate back onto the magnetic stand and incubate for about 1 min to separate

beads from solution. Wait for the solution to become clear

13. While the plate is on the magnetic stand, aspirate 20 μl clear solution from the plate and transfer

to a fresh plate. Do not disturb the beads. If beads are accidentally pipetted, resuspend

them back, wait for the solution to clear up, and repeat.

14. Proceed to checking the library integrity with gel electrophoresis or freeze the libraries.

Note: Because they do not contain EDTA they are at risk of degradation and should always be thawed slowly on ice.
